# Quantitation of human enteric viruses as alternative indicators of fecal pollution to evaluate wastewater treatment processes

**DOI:** 10.1101/2021.08.05.455335

**Authors:** Audrey Garcia, Tri Le, Paul Jankowski, Kadir Yanaç, Qiuyan Yuan, Miguel Uyaguari-Díaz

## Abstract

We investigated the potential use and quantitation of human enteric viruses in municipal wastewater samples of Winnipeg (Manitoba, Canada) as alternative indicators of contamination and evaluated the processing stages of the wastewater treatment plant. During the fall 2019 and winter 2020 seasons, samples of raw sewage, activated sludge, effluents, and biosolids (sludge cake) were collected from the North End Sewage Treatment Plant (NESTP), which is the largest wastewater treatment plant in the City of Winnipeg. DNA and RNA enteric viruses, as well as the *uidA* gene found in *Escherichia coli* were targeted in the samples collected from the NESTP. Total nucleic acids from each wastewater treatment sample were extracted using a commercial spin-column kit. Enteric viruses were quantitated in the extracted samples via quantitative PCR using TaqMan assays.

The average gene copies assessed in the raw sewage were not significantly different (p-values ranged between 0.0547 and 0.7986) than the average gene copies assessed in the effluents for Adenovirus and crAssphage (DNA viruses), Pepper Mild Mottle Virus (RNA virus), and *uidA* in terms of both volume and biomass. A significant reduction of these enteric viruses was observed consistently in activated sludge samples compared with those for raw sewage. Corresponding reductions in gene copies per volume and gene copies per biomass were also seen for *uidA* but were not statistically significant (p-value = 0.8769 and p-value = 0.6353, respectively). The higher gene copy numbers of enteric viruses and *E. coli* observed in the effluents may be associated with the 12-hour hydraulic retention time in the facility. Enteric viruses found in gene copy numbers were at least one order of magnitude higher than the *E. coli* marker *uidA*. This indicate that enteric viruses may survive the wastewater treatment process and viral-like particles are being released into the aquatic environment. Our results suggest that Adenovirus, crAssphage, and Pepper mild mottle virus can be used as complementary viral indicators of human fecal pollution.

## INTRODUCTION

The human fecal waste present in raw sewage (RS) contains pathogens that can cause numerous diseases. This can have a huge negative impact to public, aquatic health, and the economy (Stachler, et al., 2017). Wastewater treatment plants (WWTPs) serve as protective barriers between communities and the environment by reducing the organic matter present in wastewater. Water quality is currently assessed using traditional markers such as coliforms and *Escherichia coli*, leaving other microbes such as viruses largely unexplored. The North End Sewage Treatment Plant (NESTP) in Winnipeg, Manitoba handles approximately 70% of the city’s wastewater treatment, serving over 400,000 people (City of Winnipeg, Water and Waste Department, 2020). The treatment process at the NESTP first involves RS undergoing primary treatment to remove solids. During the next treatment cycle, activated sludge (AS), a heterotrophic cocktail of bacteria and protozoa, degrades organic matter present in solid waste. The activated sludge (also known as biological treatment or secondary treatment) is the most widely used process around the world to treat municipal wastewater (Racz et al., 2010; Scholz, 2016), and its use will likely continue for centuries as it is a cheap and efficient treatment process. After the biological treatment, wastewater is UV-disinfected and discharged as effluents (EF) into the river (City of Winnipeg, Water and Waste Department, 2020). Approximately 200 million liters of EF are discharged per day (City of Winnipeg, Water and Waste Department, 2020).

The main indicator of biological contamination used in wastewater treatment screening is *E. coli*, a fecal coliform bacterium (Hood et al., 1983). It is present in the gut of humans and warm-blooded animals, and widely used as the main indicator of fecal pollution during the wastewater treatment process. Nevertheless, the use of only fecal bacteria indicator in wastewater excludes other possible pathogen groups present, such as human enteric viruses. Targeting these viruses in EF could be an alternative method to monitor the wastewater treatment process. Within this context, Dutilh et al. (2014) targeted the DNA crAssphage genome in a human fecal sample. With further bioinformatics testing, it was predicted that the crAssphage genome is highly abundant, and it was identified in 73% of human fecal metagenomes surveyed (Dutilh, et al., 2014). In a study conducted by Zhang et al. (2006), the most abundant fecal virus found in dry weight fecal matter was the plant RNA virus, Pepper mild mottle virus (PMMV).

In the present study, samples of RS, AS, EF, and biosolids/sludge cake (SC) from the NESTP were collected (during fall 2019 and winter 2020) to investigate the potential of quantitating human enteric viruses in wastewater samples as complementary indicators of contamination to evaluate the processing stages of wastewater treatment. DNA enteric viruses in this study include human Adenovirus (AdV) and cross assemblied phage (crAssphage), while RNA enteric viruses include PMMV, Noroviruses (NoV) of the genogroups GI and GII, Astrovirus (AstV), Sapovirus (SaV), and Rotavirus (RoV). We also studied the presence of a molecular marker for *E. coli*, the *uidA* gene, in the samples collected from the NESTP. An overview of the workflow is illustrated in *Fig. 1*.

**Figure 1.**
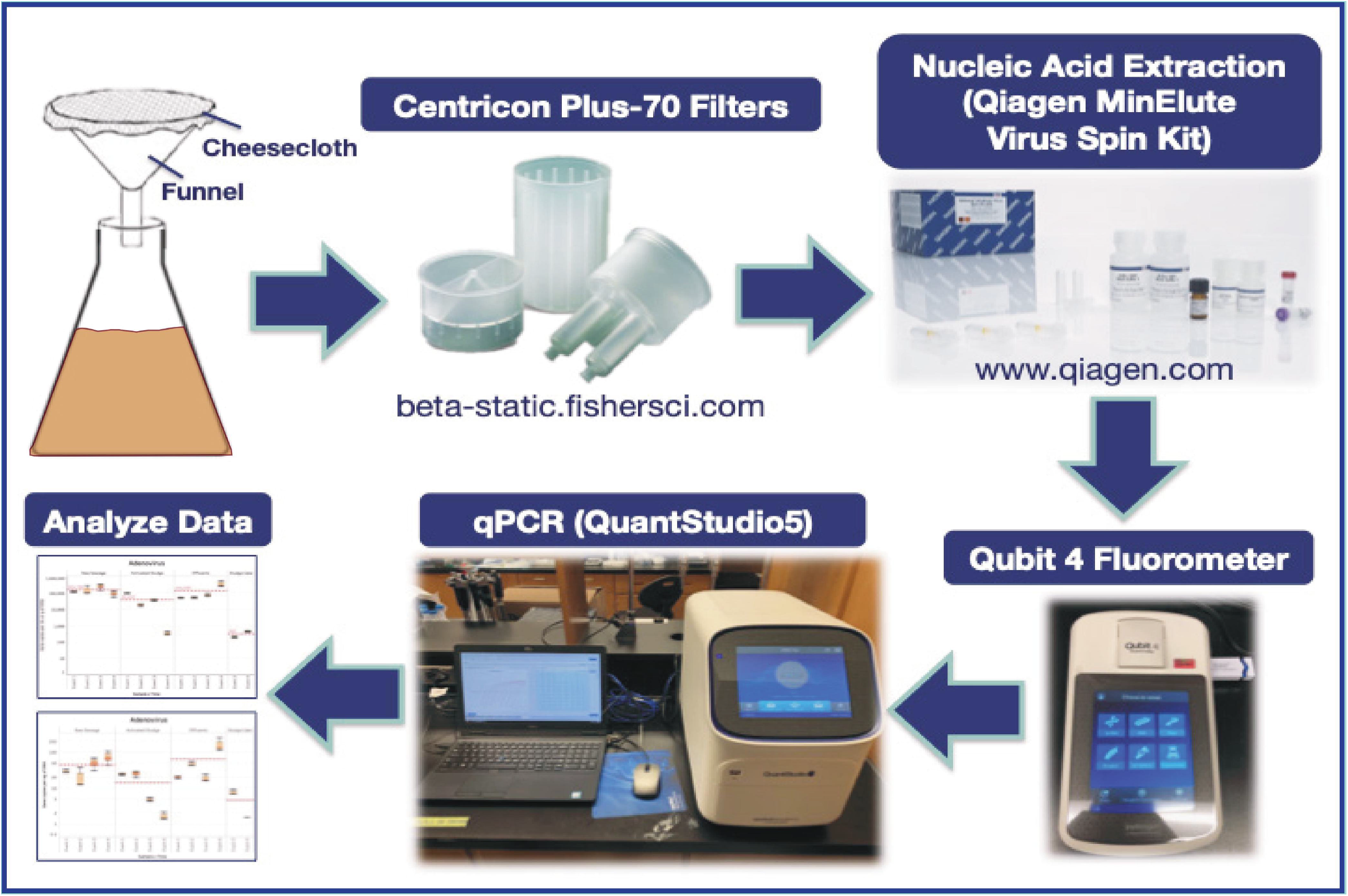
Graphical abstract of workflow.

## MATERIALS AND METHODS

### Sample Collection

A liter of RS, AS, EF, and 1 kg of SC were collected from the NESTP during each of the sampling events in fall 2019 and winter 2020. Each sample was sealed in a 1-L sterile polyethylene container lined with a sterile plastic bag. Samples were collected on October 22^nd^, 2019 (Event 1) and November 28^th^, 2019 (Event 2) in the fall season. In the winter season, samples were collected on December 18^th^, 2019 (Event 3) and February 6^th^, 2020 (Event 4). SC samples were collected earlier in the day during Events 3 and 4. All samples were kept at 4°C and processed within 24 hours of collection.

### Assessment of Ultrafiltration for Viral Recovery Efficiency

Armored RNA (Asuragen, Inc., Austin, TX, USA), an artificial virus, was used to assess recovery efficiency of the ultrafiltration method employed herein. We spiked in 40,000 copies of Armored RNA into 120 mL raw sewage samples collected in duplicates from the NESTP, but this was not included as part of this study. Primers (381F: 5’-AGCCTGTCAATACCTGCACC-3’ and 475R: 5’-CACGCTTAGATCTCCGTGCT-3’), and probe (420P: 5’ Cy5-AGAGTATGAGAGGTCGACGA-TAO 3’) were designed using Primer design tool of Geneious Prime version 2021.1.1 (https://www.geneious.com) and targeted a 95-bp region within the Armored RNA genome. This targeted 95-bp fragment was sent to Integrated DNA Technologies (IDT, Inc., Coralville, Iowa) to synthetize a gBlock construct. Serial dilutions of this synthetic fragment were used to generate standards and quantify gene copy numbers (GCNs) of Armored RNA via quantitative reverse transcription PCR (RT-qPCR). Standard and raw sewage samples were run in triplicates.

Thermal cycling reactions were performed at 50°C for 5 minutes, followed by 45 cycles at 95°C for 10 seconds and 60°C for 30 seconds on a QuantStudio 5 Real-Time PCR System (Life Technologies, Carlsbad, CA, USA). Each 10-μl RT-qPCR mixture consisted of 2.5 µL 4X TaqMan Fast Virus 1-Step Master Mix (Life Technologies, Carlsbad, CA, USA), 400 nM each primer, 200 nM probe, and 2.5 μl of template and ultrapure DNAse/RNAse free distilled water (Promega Corporation, Fitchburg, WI, USA).

### Ultrafiltration of Wastewater Samples

Each wastewater treatment sample (RS, AS, and EF), including Millipore Milli-Q water as a negative control, was first filtered via a funnel and cheesecloth to remove any solid waste or debris. Next, 140 mL of each wastewater sample was concentrated using an ultrafiltration method with Centricon Plus-70 filter units (Millipore Corporation, Billerica, MA, USA). The ultrafiltration process used a sterile glass pipette, where 70 mL of each wastewater sample was added into their correspondingly labeled sample filter cup pre-assembled with the filtrate collection cup. Each assembly was then sealed with a cap. The Centricon Plus-70 assemblies were placed into a swinging bucket rotor and centrifuged at 3000 × g for 30 minutes at 20°C. Subsequently, the filtrate was discarded, and the remaining 70 mL of the samples was added into their correspondingly labeled sample filter cup pre-assembled with the filtrate collection cup. Samples were spun at the same speed and temperature for 45 minutes. After centrifugation, the sample filter cup was separated from the filtrate collection cup. The concentration collection cup was then turned upside down and placed on top of the sample filter cup. The device was carefully inverted and placed into the centrifuge. Centricon Plus-70 filter units were centrifuged at 800 × g for 2 minutes at 20°C. After this step, the concentrated sample was collected from the concentration cup via a micropipette. The final volume was measured for each wastewater sample. If needed, 10 mM Tris-HCl, pH 8.5 buffer (Qiagen Sciences, Maryland, MD) was added to the concentrate to make up a total volume of 250 μL. If the final volume of the concentrate was over 250 μL, Tris buffer was not added. Aliquots containing 250 μL were made and stored at 4°C and processed within 24h.

### Sludge Cake Preparation for Ultrafiltration

To remove cells from the SC samples, a 1X phosphate-buffered solution (PBS) with 0.15M NaCl, 0.05% Tween-20, and pH 7.5 was used. Approximately 30 g of SC sample per sampling event (Events 3 and 4) was collected and divided into six Falcon tubes for each event (∼5-6 g per tube). Approximately 30 mL of PBS was added to each tube. The Falcon tubes filled with SC samples were homogenized at constant agitation for 15 minutes at 2500 rpm in a vortex mixer. These tubes were then centrifuged at a speed of 4500 × g for 50 minutes. The supernatant from each tube was subsequently recovered and transferred to a new sterile Falcon tube. For each sample event, 140 mL of supernatant was used for ultrafiltration as described previously.

### Nucleic Acid (DNA/RNA) Extraction and Fluorometric Assessment

Once the final volume of concentrate was collected from each wastewater sample, the sample was pretreated with InhibitEX buffer (Qiagen Sciences, Maryland, MD) as indicated by the manufacturer. Then, QIAamp MinElute virus spin kit (Qiagen Sciences, Maryland, MD) was used to extract total nucleic acids from each wastewater sample as per the manufacturer’s instructions, which included the use of Qiagen Protease and carrier RNA (Qiagen Sciences, Maryland, MD). Samples were eluted in 75 μL of Buffer AVE (Qiagen Sciences, Maryland, MD), quantified, and stored at -80°C for downstream processes. The nucleic acid concentration and purity were assessed using Qubit dsDNA high sensitivity and RNA assay kits in a Qubit 4 fluorometer (Invitrogen, Carlsbad, CA, USA), respectively. Qubit results can be found in Supplementary Materials (*Table S1*).

### qPCR Primers, Probes, and gBlocks Gene Fragments

*Table 1* summarizes the primers and probes used in this study. Forward and reverse primers listed in *Table 1* were used in the Primer-BLAST tool to extract gene target regions (Ye, et al., 2012). Extracted regions were then uploaded to the Geneious software to verify oligonucleotide sequences associated to the flanking regions and probe. The generated sequences were sent to Integrated DNA Technologies (IDT, Inc., Coralville, Iowa, USA) to generate the desired gBlocks constructs. IDT manufactured all the primers used for qPCR, as well as the probes Ast-P, Ring1a.2, and Ring 2.2 (*Table 1)*. However, probes Sav124TP, Sav5TP, NSP3-P, AdV-P, PMMV-P, and CrAss-P were manufactured by Life Technologies (Carlsbad, CA, USA).

**Table 1.**
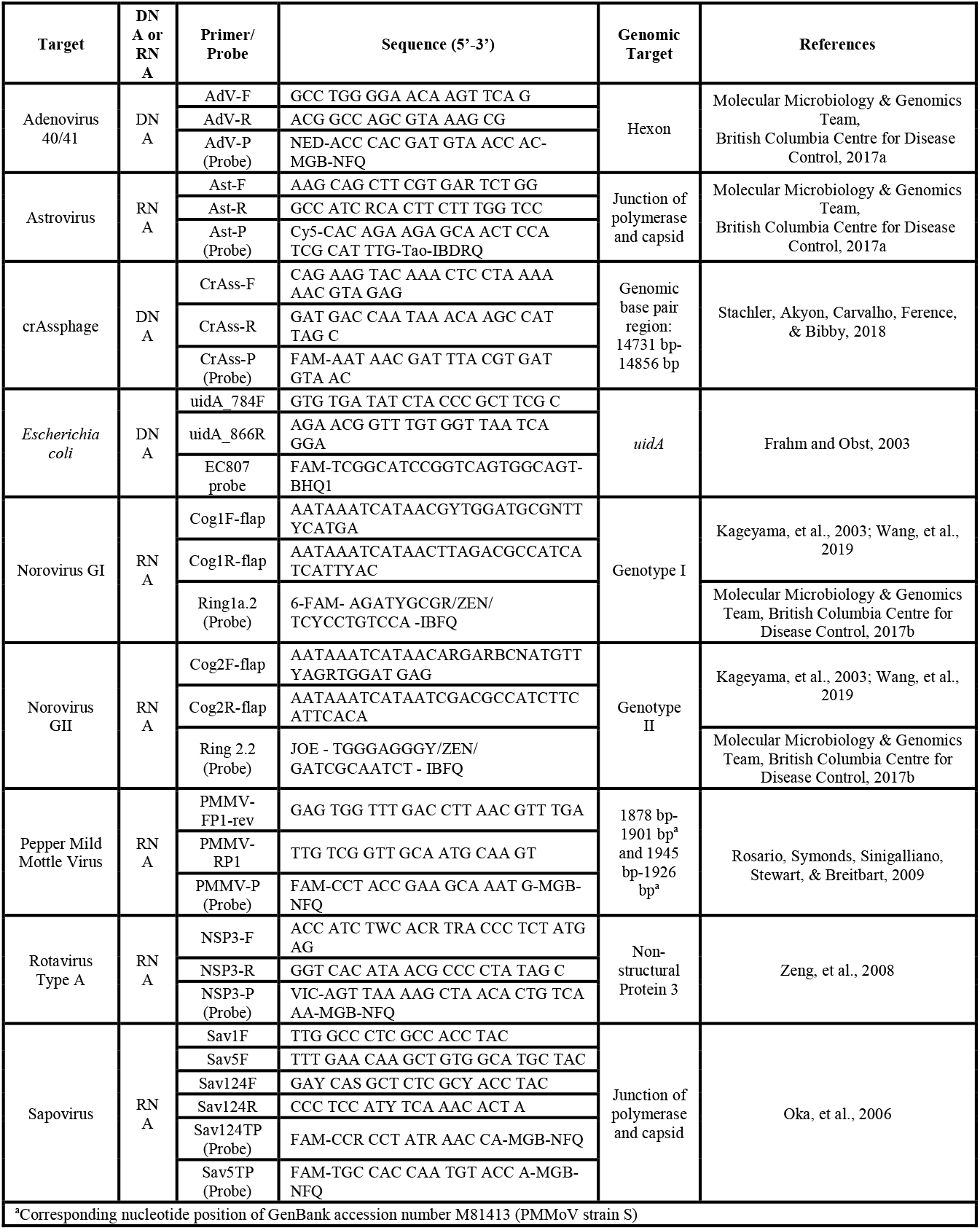
Primers and probes used in the present study.

### Quantitative PCR Assays

Taqman Environmental Master Mix 2.0 (Life Technologies, Carlsbad, CA, USA) was used for assays involving DNA enteric viruses and *uidA*, while 4x Taqman Fast Virus 1-Step Master Mix (Life Technologies, Carlsbad, CA, USA) was used for RNA enteric viruses. Each 10 μl qPCR reaction contained 500 nM of each of the forward primer and the reverse primer and 250 nM of its designated probe when targeting both DNA and RNA viruses. Five μl of Environmental Master Mix was utilized in each qPCR reaction for targeting DNA viruses, while 2.5 μl of 4x Fast Virus Master Mix was used in each qPCR reaction for targeting RNA viruses. The *uidA* qPCR reaction consisted of 5 μl of Environmental Master Mix, 400 nM of each primer, and 100 nM of probe. All qPCR reactions used 2 μl of template. Each qPCR reaction was performed in triplicates on an ABI QuantStudio 5 PCR system (Applied Biosystems, Foster City, CA, USA). The DNA enteric viruses (AdV and crAssphage) and *uidA* were subjected to the following conditions: 50.0°C for 2 minutes and 95.0°C for 10 minutes followed by 40 cycles of 95.0°C for 15 seconds and 60.0°C for 1 minute. The RNA enteric viruses (SaV, RoV, AstV, GI and GII NoV, PMMV) were subjected to the following conditions: 50.0°C for 5 minutes and 95.0°C for 20 seconds followed by 40 cycles of 95.0°C for 3 seconds and 60.0°C for 30 seconds. Raw qPCR output files can be found on GitHub (https://git.io/J8VJ6).

### Assessment of Gene Copy Numbers by Volume and Biomass

Gene copy numbers (GCNs) were expressed in terms of sample (per mL or g of sample) and biomass (per ng of DNA or RNA). GCNs per mL of sample were calculated as previously described by Ritalahti et al. (2006). When calculating GCNs per mL of sample, the final volume recovered after filtering 140 mL of wastewater sample was used in the formula. For the SC samples, the mass of SC collected was used in the formula to produce results in GCNs per g of sample.

### Collection of Metadata for Sampling Events

To perform Principal Component Analysis (PCA) and Spearman’s rank correlation analysis for EF samples, metadata pertinent to the sampling events was retrieved. Water quality parameters obtained from the NESTP were combined with their October 2019 monitoring data (City of Winnipeg, Water and Waste Department, 2019) to complete some of the missing fields. For a value not found in either document, data interpolation was performed by taking an average of the corresponding values for the days before and after the sampling event. In addition, the Government of Canada’s historical weather database was utilized to obtain the mean temperature on the sampling dates and the total precipitation over three days before each sampling event (hereafter referred to as “precipitation”) (Environment and Climate Change Canada, 2021). The values for all parameters were transformed using log_10_, except for precipitation due to the presence of zero values. These variables were used with log_10_-transformed GCNs per mL sample for AdV, crAssphage, PMMV, and *uidA* (targets with quantifiable qPCR readings for all replicates across all events) as input for downstream analyses (PCA and Spearman’s rank correlation analysis).

### Data Handling, Statistical Analysis, and Data Visualization

Various applications were employed to process data at different steps of the pipeline. Input data, such as output from the qPCR instrument, was subjected to rudimentary formatting and cleaning in Microsoft Excel, which was also used to calculate GCNs per mL or g sample and per ng nucleic acid. R (R Core Team, 2021) and its integrated development environment RStudio (RStudio Team, 2021) were utilized to further process the data and perform statistical analyses and output visualizations. These operations included general linear models (and estimated pairwise differences) using the package *sasLM* version 0.6.0 (Bae, 2021), PCA (corresponding biplots were created using the package *ggbiplot* version 0.55 (Vu, 2011)), and Spearman’s correlation matrix using the package *Hmisc* version 4.5-0 (Harrell Jr., 2021). The package *reshape2* version 1.4.4 (Wickham, 2020) was used to reformat these correlation matrices to enhance compatibility with other data-handling tools. Information about other packages is provided in Supplementary Materials (*Table S2*). The R script used for analysis can be found on GitHub (https://git.io/J8VUl).

Another software involved in data visualization was Tableau. Specifically, it was used to generate boxplots for GCNs per mL or g sample and per ng nucleic acid, as well as the heatmap representing the above-mentioned Spearman correlation matrix.

For all tests, a p-value of 0.05 was assumed to be the minimum level of significance.

## RESULTS

From our assessment of the ultrafiltration method used in this study, the recovery efficiency of Armored RNA as measured by RT-qPCR was estimated to be between 7.14% and 8.64% for both raw sewage samples.

The GCN values for the DNA and RNA viruses and *uidA* were transformed into log_10_ form. These values were run through a general linear model Tukey-Kramer analysis, and the means of each wastewater processing stage for each target were analyzed. The GCNs were expressed in terms of volume (mL) or weight (g) of sample and biomass (ng of nucleic acids). The presence of DNA and RNA viral gene copies and *uidA* in the Milli-Q water (negative control) samples across all Events 1-4 were negative. The orange-dotted lines in *Figs. 2-6* indicate the mean of the number of gene copies of each wastewater treatment sample across all events.

**Figure 2.**
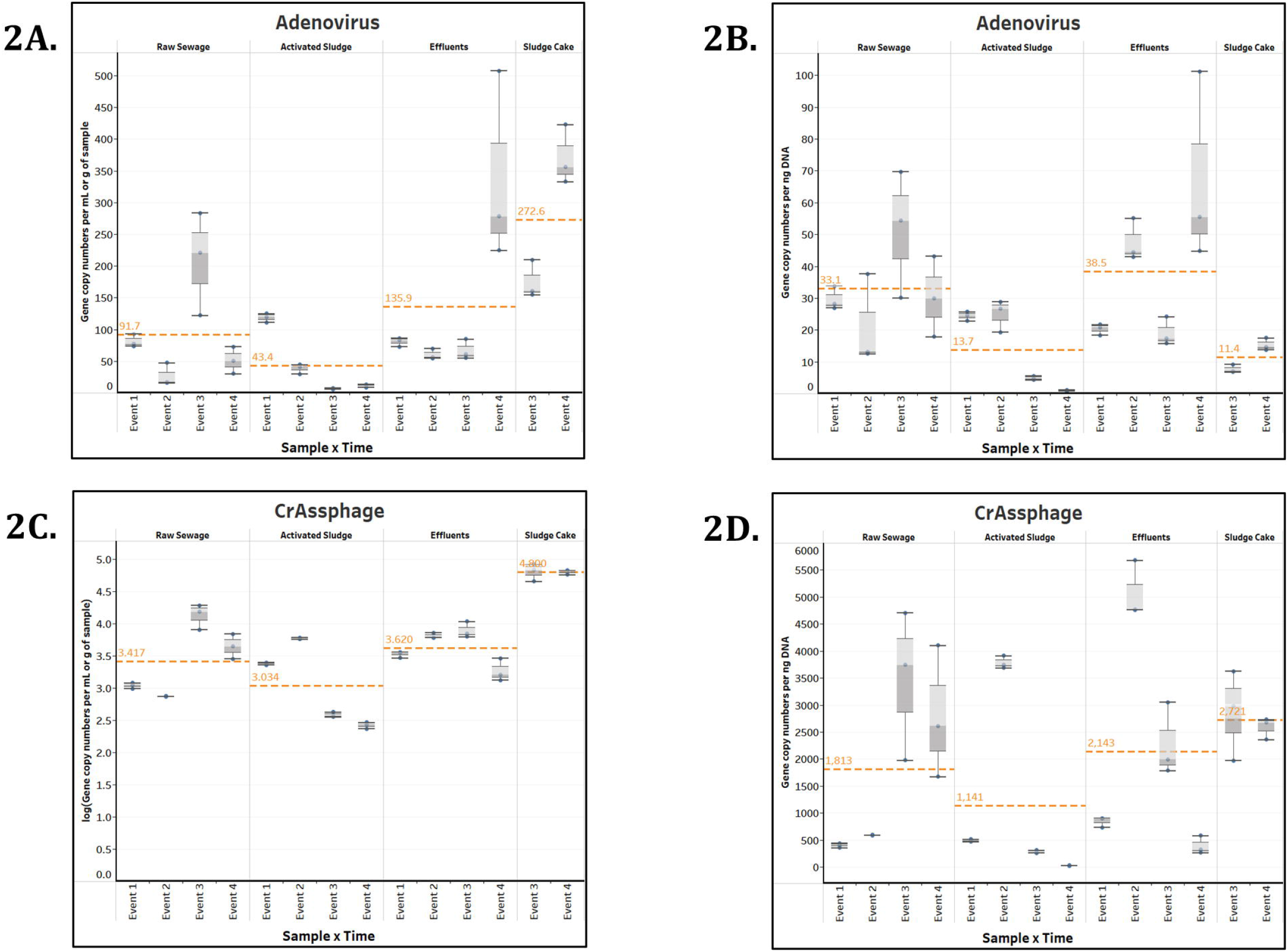
Box plots of the number of gene copies of DNA enteric viruses across each wastewater stage throughout Events 1-4. The unit for the SC in Figures 2A and 2C is gene copies per g of sample.

The average GCNs assessed in the RS were not significantly different (p-values ranged between 0.0547 and 0.7986) compared to the average GCNs assessed in the EF for the DNA enteric viruses (AdV and crAssphage) (*Fig. 2*), PMMV (*Fig. 3*), and *uidA* (*E. coli*) (*Fig. 4*) in terms of both volume and biomass. However, the average GCNs of the DNA enteric viruses assessed in AS were significantly and consistently lower compared to RS. Corresponding reductions in gene copies per volume and gene copies per biomass were also seen for *uidA*, although these reductions were not statistically significant, with p-values being 0.8769 and 0.6353, respectively. For all the aforementioned targets, there was a relatively higher number of gene copies observed in the EF across all events compared to AS samples.

**Figure 3.**
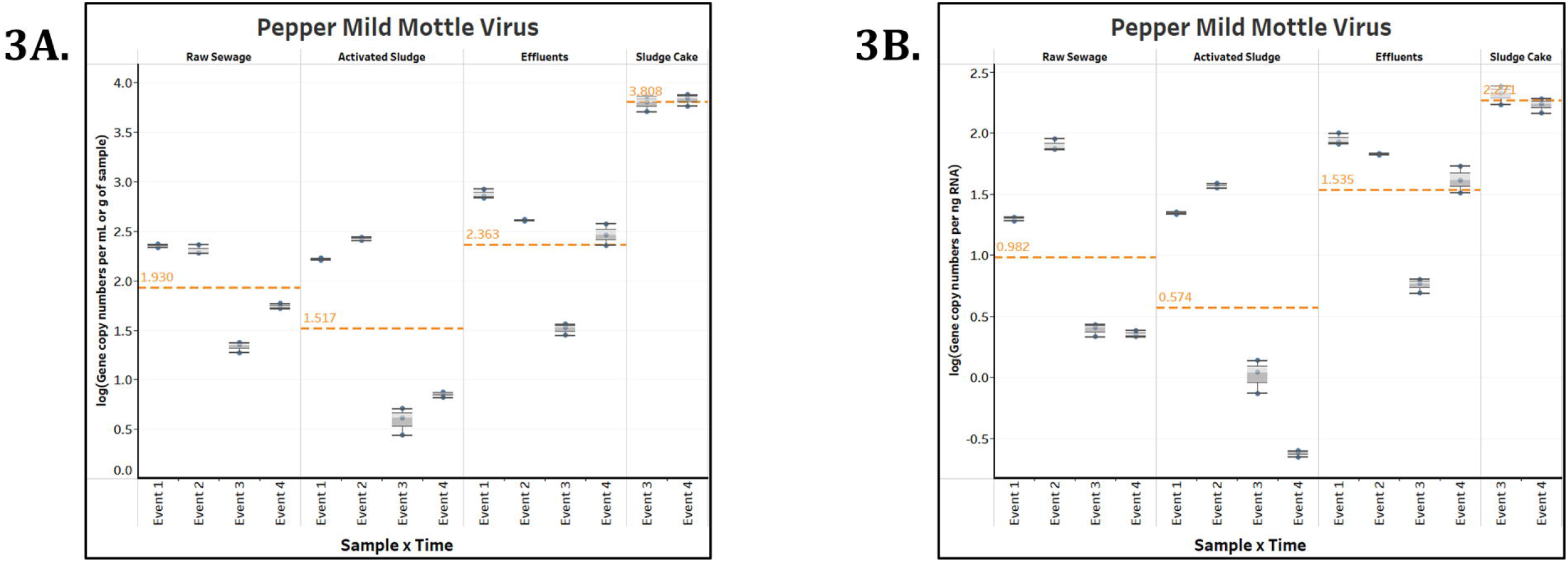
Box plots of the number of genes copies of PMMV across each wastewater stage throughout Events 1-4. The unit for the SC in Figure 3A is gene copies per g of sample.

**Figure 4.**
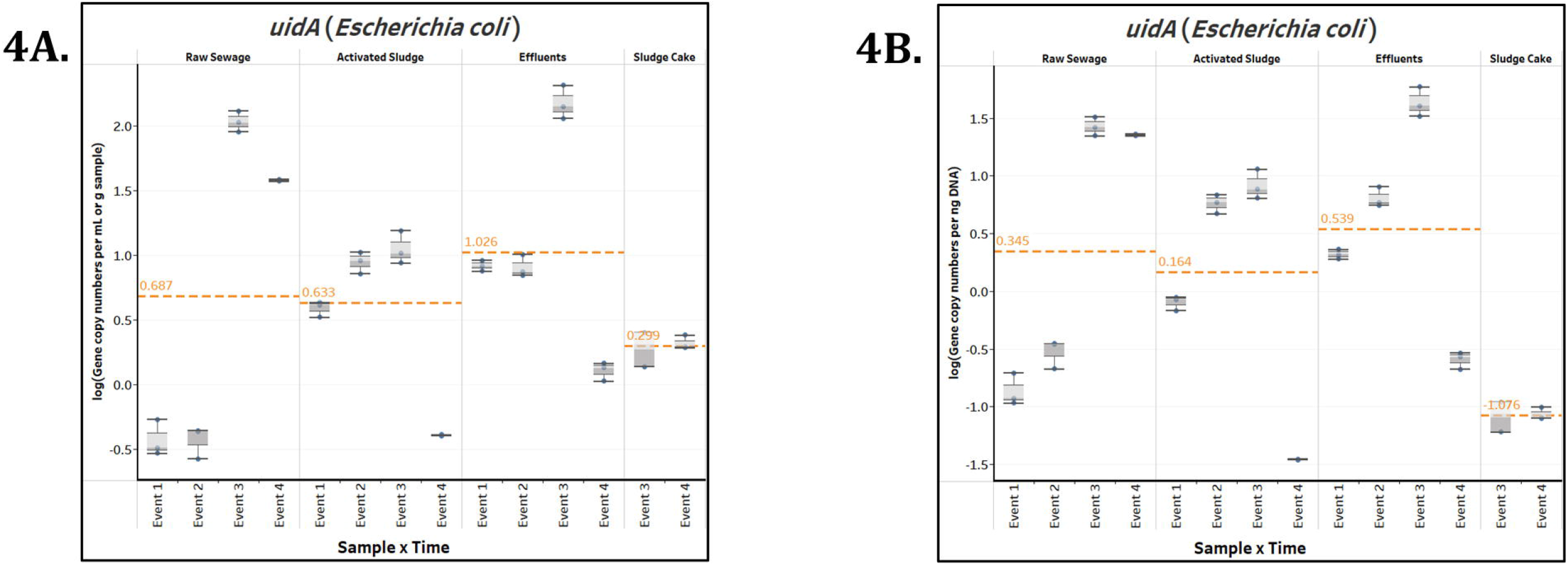
Box plots of the number of gene copies of *uidA* across each wastewater stage throughout Events 1-4. The unit for the SC in Figure 4A is gene copies per g of sample.

NoV GI and GII were also targets for our study. Boxplots of their GCNs across the different wastewater stages throughout Events 1-4 can be found in Supplementary Materials (*Fig. S1*). In Events 1 and 2 (Fall season), NoV GI was below qPCR detection limits for all samples (RS, AS, and EF). In addition, NoV GII GCNs for all samples collected in Event 2 and AS samples in Events 3 and 4 (Winter season) were also below the detection limits. Among the quantifiable samples, statistically significant GCN differences in terms of volume/mass and biomass were calculated for the pairs of AS-EF (p-values were 4.483 × 10^−6^ and 3.226 × 10^−7^ respectively), AS-RS (1.658 × 10^−6^, 1.091 × 10^−5^), and AS-SC (1.481 × 10^−9^, 4.083 × 10^−7^). No significant differences were detected among treatments for GCNs of NoV GI. There was no significant difference between the mean gene copies of NoV GII in the RS and EF samples in terms of volume (p-value = 0.7377), but the difference was significant in terms of biomass (p-value = 0.04905). The corresponding quantities of all the other sample pairs were statistically significant when looking at both the volume/mass and biomass perspectives, with p-values ranging from 1.304 × 10^−8^ to 0.0046, except for AS-RS GCN difference in terms of biomass (p-value = 0.0637).

RoV gene copies across the various wastewater treatment stages from Event 1 to 4 were also examined. The boxplots illustrating these results in terms of both sample and biomass can be found in the Supplementary Materials (*Fig. S2*). RoV GCNs were below detection limit for all samples collected in Events 1 and 2. Looking at the EF-SC pair, the mean GCNs differed significantly in terms of volume/mass (p-value = 2.649 × 10^−7^) but not biomass (p-value = 0.4298). No significant GCN differences could be detected between RS and AS samples in terms of both volume (p-value = 0.4155) and biomass (p-value = 0.6662). The equivalent magnitudes for the remaining pairs per volume/mass and per biomass were statistically significant, with p-values being between 7.907 × 10^−10^ and 0.02433, respectively.

In the present study, there was no detection of gene copies for AstV and SaV (Sav1, Sav124, and Sav5) in any of the wastewater samples across all events. In addition, to eliminate the possibility of inhibitors or contaminants such as humic acids, additional qPCR tests using bovine serum albumin (data not shown) were conducted with environmental samples (including AS). No significant differences were observed between samples with and without the enzyme. To investigate any potential relationship between collected data for EF samples, PCA was performed with log_10_-transformed variables. We found that three components (PC1, PC2, and PC3) explained 99.14% of the variance between variables. A summary of the weight of components is included in the Supplementary Materials (*Table S3*). PC1 and PC2 were used to create the biplot in *Fig. 5*. Biplots for PC1 versus PC3 (*Fig. S3*) and PC2 versus PC3 (*Fig. S4*) are included in the Supplementary Materials.

**Figure 5.**
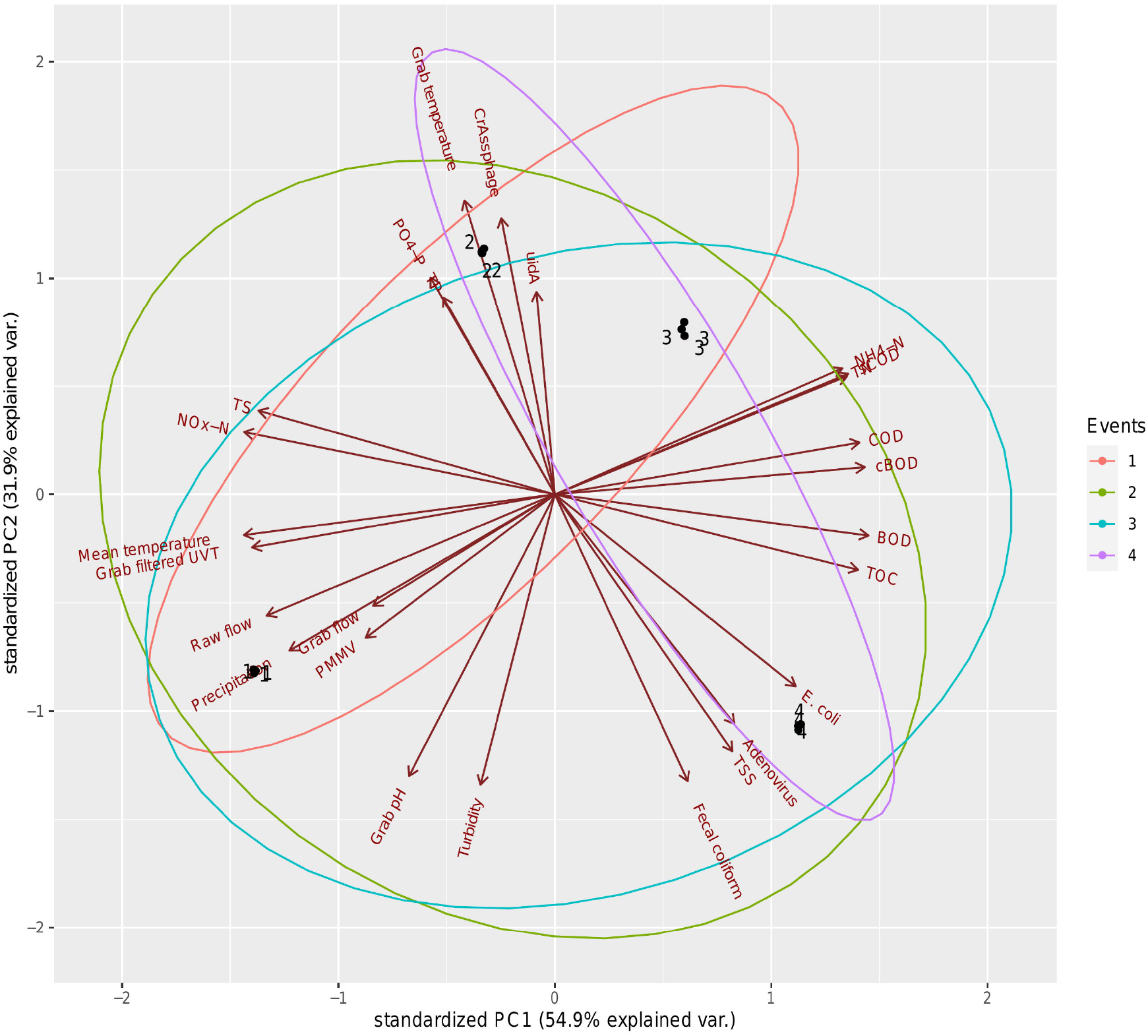
Principal Component Analysis of log_10_-transformed EF parameters, PC1 versus PC2. The only variable not log_10_-transformed was precipitation due to presence of zero values.

Overall, based on the biplot of PC1 and PC2, samples from the four events were distinct from one another, as point clusters of the four events can be seen occupying different quadrants. PC1, explaining 54.9% of the observed variance, received a notable and positive contribution from COD, cBOD, BOD, and TOC. Strongly negative contributors to PC1 were mean temperature, grab filtered UVT, NO_x_-N, and TS. These observations were supported by subsequent Spearman’s rank correlation analysis (*Fig. 6*), as COD, cBOD, BOD, and TOC demonstrated strongly positive correlations with one another (rho ranging between 0.8000 and 0.9487) (p-value < 0.005) and strongly negative correlations with mean temperature, grab filtered UVT, NO_x_-N, and TS (rho ranging between -1.000 and -0.8000) (p-value < 0.005). PC2 explained 31.9% of the variance between sampling events and showed a strong contribution from crAssphage, *uidA*, and grab temperature. This observation was also supported by the Spearman’s rank correlation analysis showing these variables having strongly positive correlation with one another (rho ranging between 0.7169 and 0.9218) (p-value < 0.0100). Additionally, in the biplot, the axes representing *E. coli* and fecal coliform specifically pointed towards the same quadrant, which was reflected in their moderately positive Spearman’s coefficient (0.6325) (p-value = 0.0273).

**Figure 6.**
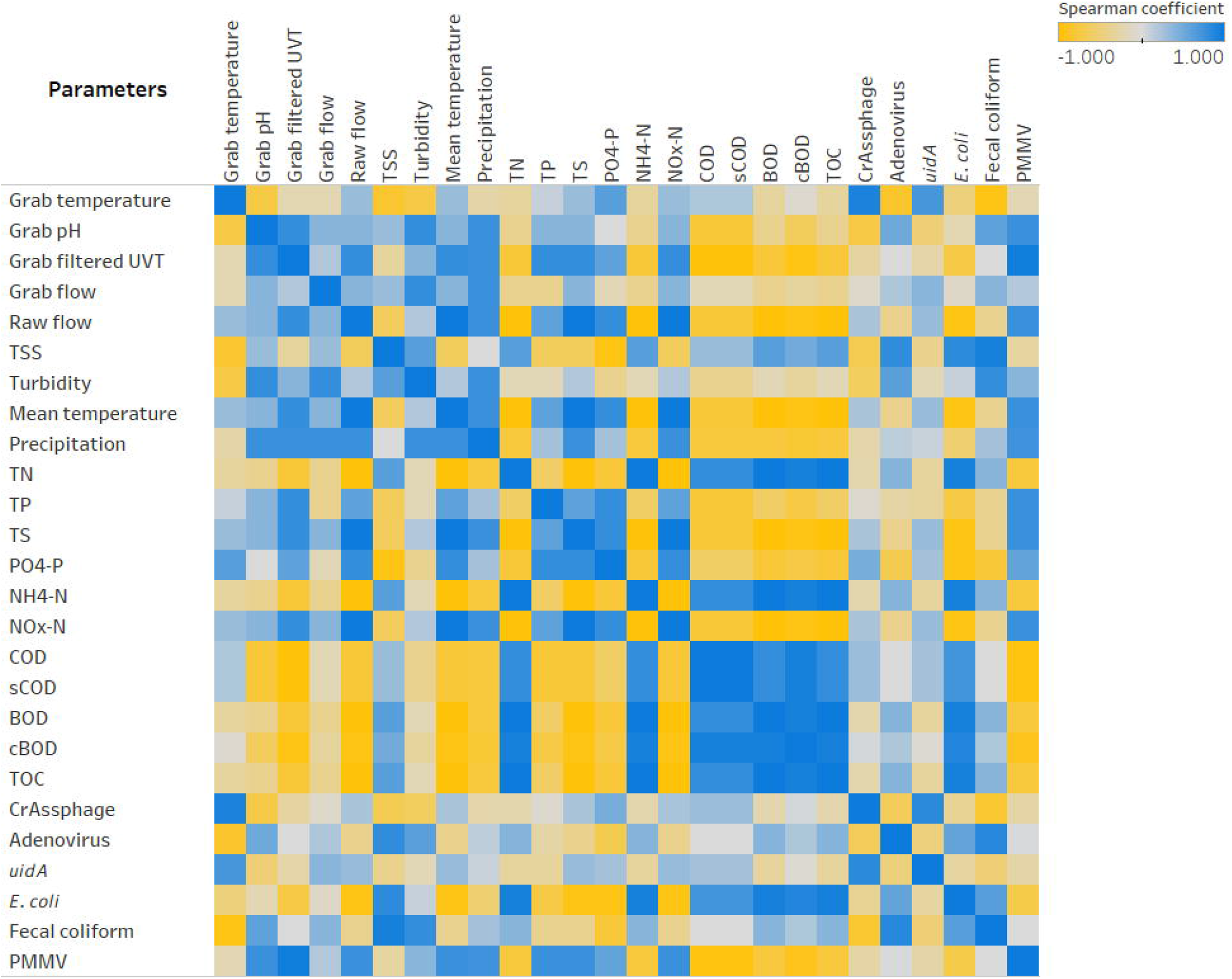
Heatmap showing Spearman’s rank correlation analysis between parameters collected for EF sampling events.

However, it is worth noting that *uidA* and *E. coli* exhibited a moderately weak negative correlation (rho = -0.3073), although it was not statistically significant (p-value = 0.3313). The two parameters with the strongest contribution against PC2 were grab pH and turbidity, which was illustrated by the strongly positive Spearman’s coefficient heatmap (rho = 0.8000) (p-value = 0.0018).

## DISCUSSION

The ultrafiltration method used in this study was assessed and the recovery efficiency was estimated to be between 7.14% and 8.64%. This range was lower compared to other methods to concentrate viral particles such as JumboSep (13.38% ± 9.11%) or skimmed milk flocculation (15.27% ± 3.32%), spiked-in wastewater samples, and using Armored RNA as internal control (Yanaç and Uyaguari, unpublished results). Viral particles may have been sorbed to biosolids present in wastewater samples. In this context, matrix has a significant effect for recovery of viral particles. When compared to other environmental matrices such as surface water samples, recovery efficiency is higher using ultrafiltration (tangential flow filtration) (32.6%±11.81%) and skimmed milk flocculation (42.64%± 15.12%) (Francis and Uyaguari, unpublished results). Samples with high concentration of particles or suspended solids tend to saturate filters and impact the recovery of viral particles (Aslan et al., 2011; Karim et al., 2009; Uyaguari-Diaz, et al., 2016).

The GCNs were expressed in terms of biomass and volume (except for SC, which was expressed in g s of sample). The higher abundance and more stable signal over time of GCNs of AdV and crAssphage (*Fig. 2*) as well as PMMV (*Fig. 3*) relative to the results of other assays make these target more representative for conducting comparisons with *E. coli*. This persistent presence is consistent with various longitudinal studies previously performed (Ballesté, et al., 2019; Farkas, et al., 2018; Farkas, et al., 2019; Hamza et al., 2019; Nour, et al., 2021; Schmitz et al., 2016; Tandukar et al., 2020; Worley□Morse et al., 2019).

A reduction of AdV, crAssphage, PMMV, and *uidA* GCNs was observed consistently in AS samples (*Figs. 2-4*). This could be a result of viral particles being sorbed to larger fractions of organic matter that had been filtered by cheesecloth early in the sample-handling process or retained in the filtration devices as previously described. It is important to mention that samples were collected within a 2-hour period from RS→AS→EF consecutively within each sampling event. The higher GCNs of viruses and *E. coli* observed in the EF may be associated with the hydraulic retention time (12 hours) in the facility and may not reflect wastewater treatment profiles at the time of collection. Other variables to consider are the overflow of sewage from rainy events and fluctuations in mixed liquor-suspended solids (Pérez et al., 2019). In our study, there were 4.6 mm of precipitation for Event 1, which may have affected the results. In the PCA analysis (*Fig. 5*), the vector for precipitation sharply denotes data points representing Event 1, indicating a possible relationship. Precipitation was also found to have positive correlations with grab flow (rho = 0.7746) and raw flow (rho = 0.7746) (*Fig. 6*). Nonetheless, further studies are needed to corroborate the potential link between precipitation and microbial counts.

Moreover, the duration of anaerobic sludge digestion is 15 days (City of Winnipeg, Water and Waste Department, 2020). In this context, GCNs of *uidA* in the SC were significantly reduced by anaerobic digestion (*Fig. 4*). This may explain why the gene copies of *uidA* in terms of biomass were lower in SC compared to all treatments (p-value < 0.02). The average gene copies across all wastewater stages (RS, AS, and EF) for *uidA* were not significantly different in terms of both volume and biomass. When compared to *uidA*, enteric viruses were found to be at least one order of magnitude more abundant than the *E. coli* marker.

GCNs of crAssphage in terms of biomass in SC were significantly higher than RS (p-value = 0.0040) and AS (p-value = 5.877 × 10^−5^) (*Fig. 2*). For PMMV, SC samples had significantly more GCNs in terms of biomass than samples from other parts of the wastewater treatment process (p-values ranged from 1.487 × 10^−5^ to 0.03788) (*Fig. 3*). Since SC is the by-product of RS and AS using anaerobic digestion, this may indicate that the presence of crAssphage and PMMV was lower in the wastewater being treated in the AS, but higher in the solids. On the other hand, GCNs of AdV in terms of biomass were not significantly different between the AS and SC samples (*Fig. 2*). Meanwhile, plant viruses such as PMMV remain more stable (in terms of biomass) during these digestion processes (Jumat, et al., 2017).

The higher presence of RoV gene copies in the EF (p-value = 0.0006592 in terms of sample and p-value = 0.001550 in terms of biomass) during the winter season (*Fig. S2*) may indicate a higher risk of transmission during cold seasons (Atabakhsh et al. 2020), since a greater presence of RoV in EF has been previously found during the winter season (Li, et al., 2011).

The negative results of SaV (Sav1, Sav124, and Sav5) across all wastewater treatment stages during the fall and winter season are consistent with Varela et al. (2018) where samples were retrieved from a wastewater treatment plant in Tunisia. Their results did not support the general belief that the peak of detection of SaV occurs during the cold and rainy months of the year. However, quantitative detection of SaV in wastewater and river water in Japan showed an increased concentration of SaV in influents between winter and spring (December to May), but a decrease in SaV concentration during the summer and autumn months (July to October) (Haramoto et al., 2008). Yet another pattern of SaV presence was reported in France, as Sima et al. (2011) found the virus to be readily detected in influents but had no clear variations in numbers over the 9-month (October to June) duration of the study. Similarly, seasonal differences in SaV concentrations were not statistically significant in a 3-year study conducted by Song et al. (2021) in China between 2017 and 2019. As a result, there are other factors that can influence wastewater SaV concentrations. For example, it has been hypothesized that isoelectric point could affect how viruses and their different strains behave in bioreactors (Miura et al., 2015). The NESTP

The gene copies of NoV GI and GII were below the detection limit in many of the AS samples (in terms of both volume and biomass), but still relatively abundant in the EF (*Fig. S1*). A possible explanation for the greatly reduced viral GCNs in AS samples is the high efficiency with which NoV GI and GII are removed, a notion supported by literature (Ibrahim et al., 2020; Kitajima et al., 2014; Schmitz et al., 2016). Furthermore, considering the observation that these viruses were found in abundance in SC samples, another contributing factor could be limitations in the sample collection process, which might not have adequately retrieved the slurry part of the sludge where the viruses are found in greater numbers as they might have sorbed to the larger fractions of the sludge solids. The relative abundance of NoV GI and GII gene copies in RS and EF during the winter months (December and February) and the absence of NoV GII in RS in October may be due to seasonal variability (Pérez, Guerrero, Orellana, Figuerola, & Erijman, 2019). However, the presence of NoV GI and GII gene copies in RS during Event 4 (February) is consistent with a study conducted by Flannery et al. (2012), in which the concentration of NoV GI and GII gene copies in the influents of a wastewater treatment plant were significantly higher during the winter months (January to March). This seasonal trend is also reflected colloquially through the virus’s sobriquet, the winter vomiting bug (Farkas, et al., 2021).

In a study conducted by El-Senousy et al. (2007), high numbers of AstV gene copies (per liter) in sewage water samples (from the Greater Cairo area in Egypt) were observed at the end of autumn and during the winter months, but the AstV concentrations tended to decrease as temperatures increased. These results are different from our findings where there was no detection of AstV in any of the wastewater treatment stages across all events. These results may be due to seasonal variability (Pérez et al., 2019) and/or reflect the pattern of infection (Corpuz et al., 2020) within the community under study.

Additionally, PCA (*Fig. 5*) and Spearman’s rank correlation analysis (*Fig. 6*) were conducted for EF samples to investigate potential connections between various physical, chemical, and biological parameters. PCA revealed that samples collected during different events from October to February were distinct from one another. This could indicate a seasonal variation in wastewater, at least in effluents. This outcome is consistent with previous literature (Comber et al., 2020). Organic chemical parameters such as COD, cBOD, BOD, and TOC were notable positive contributors to PC1, while mean temperature, grab filtered UVT, NO_x_-N, and TS most negatively contributed to PC1. These observations were validated by subsequent Spearman’s rank correlation analysis showing statistically significant coefficients. Grab filtered UVT being inversely correlated with COD, cBOD, BOD, and TOC is consistent with the widespread use of UV radiation to regulate microbial growth in a variety of medium, including water (Raeiszadeh & Adeli, 2020). Furthermore, it had been suggested that UV is an important influence to the survival of pathogens in wastewater environments, especially in cold weather conditions, such as those found in Manitoba during the surveying period (Murphy, 2017). The NESTP uses UV disinfection. Further studies are needed to evaluate the survival of enteric viruses in these reservoirs by using modification to the biological treatment and/or the disinfection process. Some of these modifications include fixed bed reactors (Sizirici & Yildiz, 2020), biofilm systems such as membrane bioreactors, biofilters, biofiltration, and carriers (Zhao et al., 2019). Other disinfection processes include the use of chlorine (liquid sodium hypochlorite solution, solid calcium hypochlorite) or newer methods such as ozone (Mezzanotte et al., 2007; Abou-Elela et al., 2012; Collivignarelli et al., 2018).

There is a possibility that viral GCNs quantified in the EF may represent an overestimation of the actual number of infectious viral particles since qPCR detects both infective and non-infective agents and UV treatment influences viral viability (Lizasoain et al., 2017). Thus, the interpretation of these results must be performed with cautiousness. On the other hand, it is also possible that the non-enveloped enteric viruses (Corpuz, et al., 2020) studied here survived the wastewater treatment process. Non-enveloped viruses are more resilient than their enveloped counterparts in numerous environmental conditions and water treatment processes (La Rosa et al., 2020). This is because of the latter group’s envelope, which contains receptors needed for infection; if the envelope is lysed, infection is not possible (La Rosa et al., 2020). Various publications have noted the resilience of non-enveloped viruses after wastewater treatment (Adefisoye et al., 2016; Campos & Lees, 2014; Farkas, et al., 2019; Fitzgerald, 2015; Fong et al., 2010; Li, et al., 2021; Prevost, et al., 2015; Ruggeri, et al., 2015; Varela, et al., 2018). In this context, we have detected GCNs of AdV, crAssphage, and PMMV in environmental surface waters receiving discharges from the NESTP, two other WWTPs, and other areas radiating away from the WWTPs within the city of Winnipeg (Francis and Uyaguari, unpublished data). Therefore, despite potential factors affecting interpretation, our results still reflect the presence of several non-enveloped enteric viruses in EF samples with reasonable quantitative accuracy.

## CONCLUSION

Our study’s primary goal was to identify human enteric viruses with the potential to become alternative indicators of fecal pollution. Towards that end, we propose AdV, crAssphage, and PMMV as more stable viral indicators of water quality due to their quantifiability illustrated in this investigation and the literature. Regular monitoring of these organisms can be useful complements to current methods for assessing wastewater treatment processes. Such vigilance could be a helpful tool to assist public health efforts in the event of a viral outbreak.

Additionally, our study indicated that enteric viruses may have survived the wastewater treatment process and viral-like particles are possibly being released into the aquatic environment. Therefore, in addition to such methods as UV radiation (which is currently used in the NESTP and was shown in our study to be inversely correlated with biological parameters), we also suggest that WWTPs consider implementing modifications and/or additions (disinfection processes) to their workflow to reduce the number of viral particles across different stages of the wastewater treatment process.

## ABBREVIATIONS

AdV: Adenovirus
AS: activated sludge
AstV: Astrovirus
BOD: biochemical oxygen demand
cBOD: carbonaceous biochemical oxygen demand
COD: chemical oxygen demand
EF: effluents
GCN: gene copy number
NESTP: North End Sewage Treatment Plant
NH_4_-N: ammonium-nitrogen
NoV: Norovirus
NO_x_-N: nitrogen oxides – nitrogen
PCA: Principal Component Analysis
PMMV: Pepper mild mottle virus
PO_4_-P: orthophosphate as phosphorus
RoV: Rotavirus
RS: raw sewage
RT-qPCR: quantitative reverse transcription PCR
SaV: Sapovirus
SC: sludge cake
sCOD: soluble chemical oxygen demand
TN: total nitrogen
TOC: total organic carbon
TP: total phosphorus
TS: total solids
TSS: total suspended solids
*uidA*: β-d-glucuronidase gene
WWTP: wastewater treatment plant

## AUTHOR CONTRIBUTIONS

AG performed the experiments, analyzed the data, prepared the figures and tables, and wrote and reviewed the drafts of the manuscript.

TL analyzed the data, prepared the figures and tables, and wrote and reviewed the drafts of the manuscript.

PJ performed the experiments and reviewed the drafts of the manuscript.

KY performed the validation experiments here described and reviewed the drafts of the manuscript.

QY contributed the analysis tools and reviewed the drafts of the manuscript.

MUD designed the experiments, provided lead guidance during the experiments and analyses, contributed the analysis tools, and reviewed the drafts of the manuscript.

All authors read and approved of the final manuscript.

## ACKNOWLEDGEMENTS

Special thanks to the City of Winnipeg and Palwinder Singh, graduate student, Department of Civil Engineering at the University of Manitoba (UoM) for sample collection. Research start-up funds grant No. 322388 were assigned to Miguel Uyaguari-Diaz at the UoM. We acknowledge The Faculty of Science, UoM, collaborative grant No. 52622 (Drs. Uyaguari and Yuan). This research was conducted at the University of Manitoba. “*The University of Manitoba campuses are located on original lands of Anishinaabeg, Cree, Oji-Cree, Dakota, and Dene peoples, and on the homeland of the Métis Nation*”.

---

**Table S1. Qubit results of extracted nucleic acid samples**.

**Table S2. R packages used that were not mentioned in the manuscript**.

**Table S3. Summary of weight of components of PCA for EF samples and related metadata**.

**Figure S1. Box plots of the number of gene copies of Noroviruses GI and GII across each wastewater stage throughout Events 1-4**.

The unit for the SC in Figures S1A and S1C is gene copies per g of sample.

**Figure S2. Box plots of the number of gene copies of Rotavirus across each wastewater stage throughout Events 1-4**.

The unit for the SC in Figure S2A is gene copies per g of sample.

**Figure S3. Principal Component Analysis of log**_**10**_**-transformed EF parameters, PC1 versus PC3**.

The only variable not log_10_-transformed was precipitation due to presence of zero values.

**Figure S4. Principal Component Analysis of log**_**10**_**-transformed EF parameters, PC2 versus PC3**.

The only variable not log_10_-transformed was precipitation due to presence of zero values.

## REFERENCES

Abou-Elela, S. I., El-Sayed, M. M. H., El-Gendy, A. S., & Abou-Taleb, E. M. (October 2012). Comparative study of disinfection of secondary treated wastewater using chlorine, UV and ozone. Journal of Applied Sciences Research, pp. 5190-5197 ref.12

Adefisoye, M. A., Nwodo, U. U., Green, E., & Okoh, A. I. (2016). Quantitative PCR Detection and Characterisation of Human Adenovirus, Rotavirus and Hepatitis A Virus in Discharged Effluents of Two Wastewater Treatment Facilities in the Eastern Cape, South Africa. Food and Environmental Virology, 8, 262–274. doi:10.1007/s12560-016-9246-4

Aslan, A., Xagoraraki, I., Simmons, F., Rose, J., & Dorevitch, S. (2011, August 19). Occurrence of adenovirus and other enteric viruses in limited-contact freshwater recreational areas and bathing waters. Journal of Applied Microbiology, 111(5), 1250–1261. doi:10.1111/j.1365-2672.2011.05130.x

Bae, K.-S. (2021). sasLM: ‘SAS’ Linear Model. Retrieved from https://CRAN.R-project.org/package=sasLM

Ballesté, E., Pascual-Benito, M., Martín-Díaz, J., Blanch, A. R., Lucena, F., Muniesa, M., … García-Aljaro, C. (2019, May 15). Dynamics of crAssphage as a human source tracking marker in potentially faecally polluted environments. Water Research, 155, 233–244. doi:10.1016/j.watres.2019.02.042

Campos, C. J., & Lees, D. N. (2014, June). Environmental Transmission of Human Noroviruses in Shellfish Waters. Applied and Environmental Microbiology, 80(12), 3552–3561. doi:10.1128/AEM.04188-13

City of Winnipeg, Water and Waste Department. (2019, October). North End Water Pollution Control Centre Monitoring Data. Winnipeg, Manitoba, Canada. Retrieved July 21, 2021, from https://www.winnipeg.ca/waterandwaste/pdfs/sewage/ComplianceReporting/2019/oct/newpcc.pdf

City of Winnipeg, Water and Waste Department. (2020, October 8). Sewage Treatment Plants. Retrieved July 21, 2021, from City of Winnipeg: https://www.winnipeg.ca/waterandwaste/sewage/treatmentPlant/default.stm#tab-north-end-sewage-treatment-plant.

Collivignarelli, M.C., Abbà, A., Benigna, I., Sorlini, S., & Torretta, V. (2018). Overview of the Main Disinfection Processes for Wastewater and Drinking Water Treatment Plants. Sustainability, 10, 86. https://doi.org/10.3390/su10010086.

Comber, S. D., Gardner, M. J., & Ellor, B. (2020, September). Seasonal variation of contaminant concentrations in wastewater treatment works effluents and river waters. Environmental Technology, 41(21), 2716–2730. doi:10.1080/09593330.2019.1579872

Corpuz, M. V., Buonerba, A., Vigliotta, G., Zarra, T., Ballesteros Jr, F., Campiglia, P., … Naddeo, V. (2020, November 25). Viruses in wastewater: occurrence, abundance and detection methods. Science of the Total Environment, 745. doi:10.1016/j.scitotenv.2020.140910

Dutilh, B. E., Cassman, N., McNair, K., Sanchez, S. E., Silva, G. G., Boling, L., … Edwards, R. A. (2014). A highly abundant bacteriophage discovered in the unknown sequences of human faecal metagenomes. Nature Communications, 5(4498), 1–11. doi:10.1038/ncomms5498

El-Senousy, W. M., Guix, S., Abid, I., Pintó, R. M., & Bosch, A. (2007, January). Removal of astrovirus from water and sewage treatment plants, evaluated by a competitive reverse transcription-PCR. Applied and Environmental Microbiology, 73(1), 164–7. doi:10.1128/AEM.01748-06

Environment and Climate Change Canada. (2021). Historical Data. Retrieved July 21, 2021, from Government of Canada: https://climate.weather.gc.ca/historical_data/search_historic_data_e.html

Farkas, K., Adriaenssens, E. M., Walker, D. I., McDonald, J. E., Malham, S. K., & Jones, D. L. (2019, June). Critical Evaluation of CrAssphage as a Molecular Marker for Human-Derived Wastewater Contamination in the Aquatic Environment. Food and Environmental Virology, 11(2), 113–119. doi:10.1007/s12560-019-09369-1

Farkas, K., Green, E., Rigby, D., Cross, P., Tyrrel, S., Malham, S. K., & Jones, D. L. (2021, May 27). Investigating awareness, fear and control associated with norovirus and other pathogens and pollutants using best–worst scaling. Scientific Reports, 11.

Farkas, K., Marshall, M., Cooper, D., McDonald, J. E., Malham, S. K., Peters, D. E., … Jones, D. L. (2018, November). Seasonal and diurnal surveillance of treated and untreated wastewater for human enteric viruses. Environmental Science and Pollution Research, 25(33), 33391–33401. doi:10.1007/s11356-018-3261-y

Fitzgerald, A. (2015). Review of Approaches for Establishing Exclusion Zones for Shellfish Harvesting around Sewage Discharge Points - Desk Study to Inform Consideration of the Possible Introduction of Exclusion Zones as a Control for Norovirus in Oysters.

Technical Report, Aquatic Water Services Ltd. Retrieved July 26, 2021, from https://webarchive.nationalarchives.gov.uk/20150418173120/ http://www.food.gov.uk/sites/default/files/Exclusion%20Zones%20Project%20FS513404%20-%20Technical%20Report%20FINAL.pdf

Fong, T.-T., Phanikumar, M. S., Xagoraraki, I., & Rose, J. B. (2010, February). Quantitative detection of human adenoviruses in wastewater and combined sewer overflows influencing a Michigan river. Applied and Environmental Microbiology, 76(3), 715–23. doi:10.1128/AEM.01316-09

Frahm, E., & Obst, U. (2003). Application of the fluorogenic probe technique (TaqMan PCR) to the detection of Enterococcus spp. and Escherichia coli in water samples. Journal of Microbiological Methods, 52(1), 123–31. doi:10.1016/s0167-7012(02)00150-1

Genz, A., Bretz, F., Miwa, T., Mi, X., Leisch, F., Scheipl, F., & Hothorn, T. (2021). mvtnorm: Multivariate Normal and t Distributions. Retrieved from http://CRAN.R-project.org/package=mvtnorm

Hamza, H., Rizk, N. M., Gad, M. A., & Hamza, I. A. (2019, November). Pepper mild mottle virus in wastewater in Egypt: a potential indicator of wastewater pollution and the efficiency of the treatment process. Archives of Virology, 164(11), 2707–2713. doi:10.1007/s00705-019-04383-x

Haramoto, E., Katayama, H., Phanuwan, C., & Ohgaki, S. (2008, March). Quantitative detection of sapoviruses in wastewater and river water in Japan. Letters in Applied Microbiology, 46(3), 408–13. doi:10.1111/j.1472-765X.2008.02330.x

Harrell Jr., F. E. (2021). Hmisc: Harrell Miscellaneous. Retrieved from https://CRAN.R-project.org/package=Hmisc

Hood, M. A., Ness, G. E., & Blake, N. J. (1983). Relationship among fecal coliforms, Escherichia coli, and Salmonella spp. in shellfish. Applied and environmental microbiology, 45(1), 122–126. doi:10.1128/aem.45.1.122-126.1983

Ibrahim, C., Hammami, S., Khelifi, N., Pothier, P., & Hassen, A. (2020). The Effectiveness of Activated Sludge Procedure and UV-C 254 in Norovirus Inactivation in a Tunisian Industrial Wastewater Treatment Plant. Food and Environmental Virology, 12, 250–259. doi:10.1007/s12560-020-09434-0

Jumat, M. R., Hasan, N. A., Subramanian, P., Heberling, C., Colwell, R. R., & Hong, P.-Y. (2017). Membrane Bioreactor-Based Wastewater Treatment Plant in Saudi Arabia: Reduction of Viral Diversity, Load, and Infectious Capacity. Water, 9 (7). doi:10.3390/w9070534

Kageyama, T., Kojima, S., Shinohara, M., Uchida, K., Fukushi, S., Hoshino, F. B., … Katayama, K. (2003, April). Broadly Reactive and Highly Sensitive Assay for Norwalk-Like Viruses Based on Real-Time Quantitative Reverse Transcription-PCR. Journal of Clinical Microbiology, 41(4), 1548–1557. doi:10.1128/JCM.41.4.1548-1557.2003

Karim, M. R., Rhodes, E. R., Brinkman, N., Wymer, L., & Fout, G. S. (2009, April). New Electropositive Filter for Concentrating Enteroviruses and Noroviruses from Large Volumes of Water. Applied and Environmental Microbiology, 75(8), 2393–2399. doi:10.1128/AEM.00922-08

Kitajima, M., Iker, B. C., Pepper, I. L., & Gerba, C. P. (2014, August 1). Relative abundance and treatment reduction of viruses during wastewater treatment processes--identification of potential viral indicators. Science of the Total Environment, 488-489, 290–296. doi:10.1016/j.scitotenv.2014.04.087

La Rosa, G., Bonadonna, L., Lucentini, L., Kenmoe, S., & Suffredini, E. (2020, July 15). Coronavirus in water environments: Occurrence, persistence and concentration methods - A scoping review. Water Research, 179. doi:10.1016/j.watres.2020.115899

Li, D., Gu, A. Z., Zeng, S.-Y., Yang, W., He, M., & Shi, H.-C. (2011, May). Monitoring and evaluation of infectious rotaviruses in various wastewater effluents and receiving waters revealed correlation and seasonal pattern of occurrences. Journal of Applied Microbiology, 110(5), 1129–37. doi:10.1111/j.1365-2672.2011.04954.x

Li, X., Cheng, Z., Dang, C., Zhang, M., Zheng, Y., & Xia, Y. (2021, July). Metagenomic and viromic data mining reveals viral threats in biologically treated domestic wastewater. Environmental Science and Ecotechnology, 7. doi:10.1016/j.ese.2021.100105

Lizasoain, A., Tort, L., García, M., Gillman, L., Alberti, A., Leite, J., … Colina, R. (2017). Human enteric viruses in a wastewater treatment plant: evaluation of activated sludge combined with UV disinfection process reveals different removal performances for viruses with different features. Letters in Applied Microbiology, 66(3), 215–221. doi:DOI: 10.1111/lam.12839

Makowski, D., Ben-Shachar, M., Patil, I., & Lüdecke, D. (2020). Automated Results Reporting as a Practical Tool to Improve Reproducibility and Methodological Best Practices Adoption. Retrieved from https://github.com/easystats/report.

Mezzanotte, V., Antonelli, M., Citterio, S., & Nurizzo, C. (2007). Wastewater disinfection alternatives: chlorine, ozone, peracetic acid, and UV light. Water Environment Research, 79(12), 2373–2379. doi:10.2175/106143007x183763.

Miura, T., Okabe, S., Nakahara, Y., & Sano, D. (2015, May 15). Removal properties of human enteric viruses in a pilot-scale membrane bioreactor (MBR) process. Water Research, 75, 282–291. doi:10.1016/j.watres.2015.02.046

Molecular Microbiology & Genomics Team, British Columbia Centre for Disease Control. (2017). Detecting Norovirus by Fast Real-Time RT-PCR. British Columbia, Canada.

Molecular Microbiology & Genomics Team, British Columbia Centre for Disease Control. (2017). Performing the GI Virus Panel by Real-Time PCR Procedure. British Columbia, Canada.

Murphy, H. (2017). Persistence of Pathogens in Sewage and Other Water Types. In J. Rose, & B. Jiménez-Cisneros (Eds.), Global Water Pathogen Project (Vol. 4). E. Lansing, MI, UNESCO. doi:10.14321/waterpathogens.51

Nour, I., Hanif, A., Zakri, A. M., Al-Ashkar, I., Alhetheel, A., & Eifan, S. (2021, April 29). Human Adenovirus Molecular Characterization in Various Water Environments and Seasonal Impacts in Riyadh, Saudi Arabia. International Journal of Environmental Research and Public Health, 18(9), 4773. doi:10.3390/ijerph18094773

Oka, T., Katayama, K., Hansman, G. S., Kageyama, T., Ogawa, S., Wu, F.-T., … Takeda, N. (2006). Detection of human sapovirus by real-time reverse transcription-polymerase chain reaction. Journal of Medical Virology, 78(10), 1347–1353. doi:10.1002/jmv.20699

Pérez, M. V., Guerrero, L. D., Orellana, E., Figuerola, E. L., & Erijman, L. (2019, July 2). Time Series Genome-Centric Analysis Unveils Bacterial Response to Operational Disturbance in Activated Sludge. mSystems, 4 (4). doi:10.1128/mSystems.00169-19

Prevost, B., Lucas, F. S., Ambert-Balay, K., Pothier, P., Moulin, L., & Wurtzer, S. (2015, October). Deciphering the Diversities of Astroviruses and Noroviruses in Wastewater Treatment Plant Effluents by a High-Throughput Sequencing Method. Applied and Environmental Microbiology, 81(20), 7215–7222. doi:10.1128/AEM.02076-15.

R Core Team. (2021). R: A language and environment for statistical computing. R Foundation for Statistical Computing, Vienna, Austria. Retrieved from https://www.R-project.org/

Racz, L., T. Datta, and R. Goel. 2010. Effect of organic carbon on ammonia oxidizing bacteria in a mixed culture. Bioresource Technology, 101 (16), 6454–60.

Raeiszadeh, M., & Adeli, B. (2020, October 14). A Critical Review on Ultraviolet Disinfection Systems against COVID-19 Outbreak: Applicability, Validation, and Safety Considerations. ACS Photonics. doi:10.1021/acsphotonics.0c01245

Ritalahti, K. M., Amos, B. K., Sung, Y., Wu, Q., Koenigsberg, S. S., & Löffler, F. E. (2006). Quantitative PCR targeting 16S rRNA and reductive dehalogenase genes simultaneously monitors multiple Dehalococcoides strains. Applied and Environmental Microbiology, 72(4), 2765–74. doi:10.1128/AEM.72.4.2765-2774.2006

Rosario, K., Symonds, E. M., Sinigalliano, C., Stewart, J., & Breitbart, M. (2009). Pepper Mild Mottle Virus as an Indicator of Fecal Pollution. Applied and Environmental Microbiology, 75(22), 7261–7267. doi:10.1128/AEM.00410-09

Rosman, N. H., Anuar, A. N., Chelliapan, S., Din, M. F., & Ujang, Z. (2014, June). Characteristics and performance of aerobic granular sludge treating rubber wastewater at different hydraulic retention time. Bioresource Technology, 161, 155–61. doi:10.1016/j.biortech.2014.03.047

RStudio Team. (2021). RStudio: Integrated Development Environment for R. Boston, MA: RStudio, PBC. Retrieved from http://www.rstudio.com/

Ruggeri, F. M., Bonomo, P., Ianiro, G., Battistone, A., Delogu, R., Germinario, C., … Fiore, L. (2015, January). Rotavirus Genotypes in Sewage Treatment Plants and in Children Hospitalized with Acute Diarrhea in Italy in 2010 and 2011. Applied and Environmental Microbiology, 81(1), 241–249. doi:10.1128/AEM.02695-14

Sarkar, D. (2008). Lattice: Multivariate Data Visualization with R. New York: Springer.

Scholz, M. Chapter 15-Activated Sludge Processes, Editor(s): Miklas Scholz, Wetlands for Water Pollution Control (Second Edition), Elsevier, Pages 91-105.

Schmitz, B. W., Kitajima, M., Campillo, M. E., Gerba, C. P., & Pepper, I. L. (2016). Virus Reduction during Advanced Bardenpho and Conventional Wastewater Treatment Processes. Environmental Science & Technology, 50(17), 9524–9532. doi:10.1021/acs.est.6b01384.

Sima, L. C., Schaeffer, J., Saux, J.-C. L., Parnaudeau, S., Elimelech, M., & Guyader, F. S. (2011, August). Calicivirus Removal in a Membrane Bioreactor Wastewater Treatment Plant. Applied and Environmental Microbiology, 77(15), 5170–5177. doi:10.1128/AEM.00583-11.

Sizirici, B., & Yildiz, I. (2020). Organic matter removal via activated sludge immobilized gravel in fixed bed reactor. E3S Web of Conferences, 191, 03006, 1-5.

Song, K., Lin, X., Liu, Y., Ji, F., Zhang, L., Chen, P., … Xu, A. (2021). Detection of Human Sapoviruses in Sewage in China by Next Generation Sequencing. Food and Environmental Virology, 13, 270–280. doi:10.1007/s12560-021-09469-x

Stachler, E., Akyon, B., Carvalho, N. A., Ference, C., & Bibby, K. (2018). Correlation of crAssphage qPCR Markers with Culturable and Molecular Indicators of Human Fecal Pollution in an Impacted Urban Watershed. Environmental Science & Technology, 52(13), 7505–7512. doi:10.1021/acs.est.8b00638

Stachler, E., Kelty, C., Sivaganesan, M., Li, X., Bibby, K., & Shanks, O. C. (2017). Quantitative CrAssphage PCR Assays for Human Fecal Pollution Measurement. Environmental Science & Technology, 51(16), 9146–9154. doi:10.1021/acs.est.7b02703

Tandukar, S., Sherchan, S. P., & Haramoto, E. (2020). Applicability of crAssphage, pepper mild mottle virus, and tobacco mosaic virus as indicators of reduction of enteric viruses during wastewater treatment. Scientific Reports, 10(3616). doi:10.1038/s41598-020-60547-9

Therneau, T. (2021). A Package for Survival Analysis in R. Retrieved from https://CRAN.R-project.org/package=survival

Ushey, K., Allaire, J., Wickham, H., & Ritchie, G. (2020). rstudioapi: Safely Access the RStudio API. Retrieved from https://CRAN.R-project.org/package=rstudioapi

Uyaguari-Diaz, M. I., Chan, M., Chaban, B. L., Croxen, M. A., Finke, J. F., Hill, J. E., … Tang, P. (2016). A comprehensive method for amplicon-based and metagenomic characterization of viruses, bacteria, and eukaryotes in freshwater samples. Microbiome, 4 (20). doi:10.1186/s40168-016-0166-1

Varela, M. F., Ouardani, I., Kato, T., Kadoya, S., Aouni, M., Sano, D., & Romalde, J. L. (2018, March 1). Sapovirus in Wastewater Treatment Plants in Tunisia: Prevalence, Removal, and Genetic Characterization. Applied and Environmental Microbiology, 84 (6). doi:10.1128/AEM.02093-17

Vu, V. Q. (2011). ggbiplot: A ggplot2 based biplot. Retrieved from http://github.com/vqv/ggbiplot

Wang, X. M., Wei, Z. M., Guo, J. M., Cai, J. M., Chang, H. M., Ge, Y. M., & Zeng, M. M. (2019, November). Norovirus Activity and Genotypes in Sporadic Acute Diarrhea in Children in Shanghai During 2014–2018. The Pediatric Infectious Disease Journal, 38(11), 1085–1089. doi:10.1097/INF.0000000000002456

Wickham, H. (2011). The Split-Apply-Combine Strategy for Data Analysis. Journal of Statistical Software, 40(1), 1–29. doi:10.18637/jss.v040.i01

Wickham, H. (2016). ggplot2: Elegant Graphics for Data Analysis. New York: Springer-Verlag.

Wickham, H. (2020, April 9). reshape2: Flexibly Reshape Data: A Reboot of the Reshape Package. Retrieved from https://cran.r-project.org/package=reshape2

Wickham, H., & Bryan, J. (2019). readxl: Read Excel Files. Retrieved from https://CRAN.R-project.org/package=readxl

Wickham, H., & Bryan, J. (2021). usethis: Automate Package and Project Setup. Retrieved from https://CRAN.R-project.org/package=usethis

Wickham, H., & Seidel, D. (2020). scales: Scale Functions for Visualization. Retrieved from https://CRAN.R-project.org/package=scales

Wickham, H., François, R., Henry, L., & Müller, K. (2021). dplyr: A Grammar of Data Manipulation. Retrieved from https://CRAN.R-project.org/package=dplyr

Wickham, H., Hester, J., & Chang, W. (2021). devtools: Tools to Make Developing R Packages Easier. Retrieved from https://CRAN.R-project.org/package=devtools

WorleyLMorse, T., Mann, M., Khunjar, W., Olabode, L., & Gonzalez, R. (2019, September). Evaluating the fate of bacterial indicators, viral indicators, and viruses in water resource recovery facilities. Water Environment Research, 91(9), 830–842. doi:10.1002/wer.1096

Ye, J., Coulouris, G., Zaretskaya, I., Cutcutache, I., Rozen, S., & Madden, T. L. (2012, June 18). Primer-BLAST: a tool to design target-specific primers for polymerase chain reaction. BMC Bioinformatics, 13 (134). doi:10.1186/1471-2105-13-134

Zeileis, A., & Croissant, Y. (2010). Extended Model Formulas in R: Multiple Parts and Multiple Responses. Journal of Statistical Software, 34(1), 1–13. doi:10.18637/jss.v034.i01

Zeng, S. Q., Halkosalo, A., Salminen, M., Szakal, E. D., Puustinen, L., & Vesikari, T. (2008). One-step quantitative RT-PCR for the detection of rotavirus in acute gastroenteritis. Journal of Virological Methods, 153(2), 238–40. doi:10.1016/j.jviromet.2008.08.004

Zhang, T., Breitbart, M., Lee, W. H., Run, J.-Q., Wei, C. L., Soh, S. W., Hibberd, M. L., Liu, E. T., Rohwer, F., & Ruan, Y. (2006, January). RNA Viral Community in Human Feces: Prevalence of Plant Pathogenic Viruses. PLOS Biology, 4 (1). doi:10.1371/journal.pbio.0040003

Zhao, Y., Liu, D., Huang, W., Yang, Y., Ji, M., Nghiem, L. D., Trinh, Q. T., Tran, N. H. (2018). Insights into biofilm carriers for biological wastewater treatment processes: Current state-of-the-art, challenges, and opportunities. Bioresource Technology, 288, 121619. doi.org/10.1016/j.biortech.2019.121619.

