## Supplementary figures and images for "Quantitation of human enteric viruses as alternative indicators of fecal pollution to evaluate wastewater treatment processes"

### Figure S1

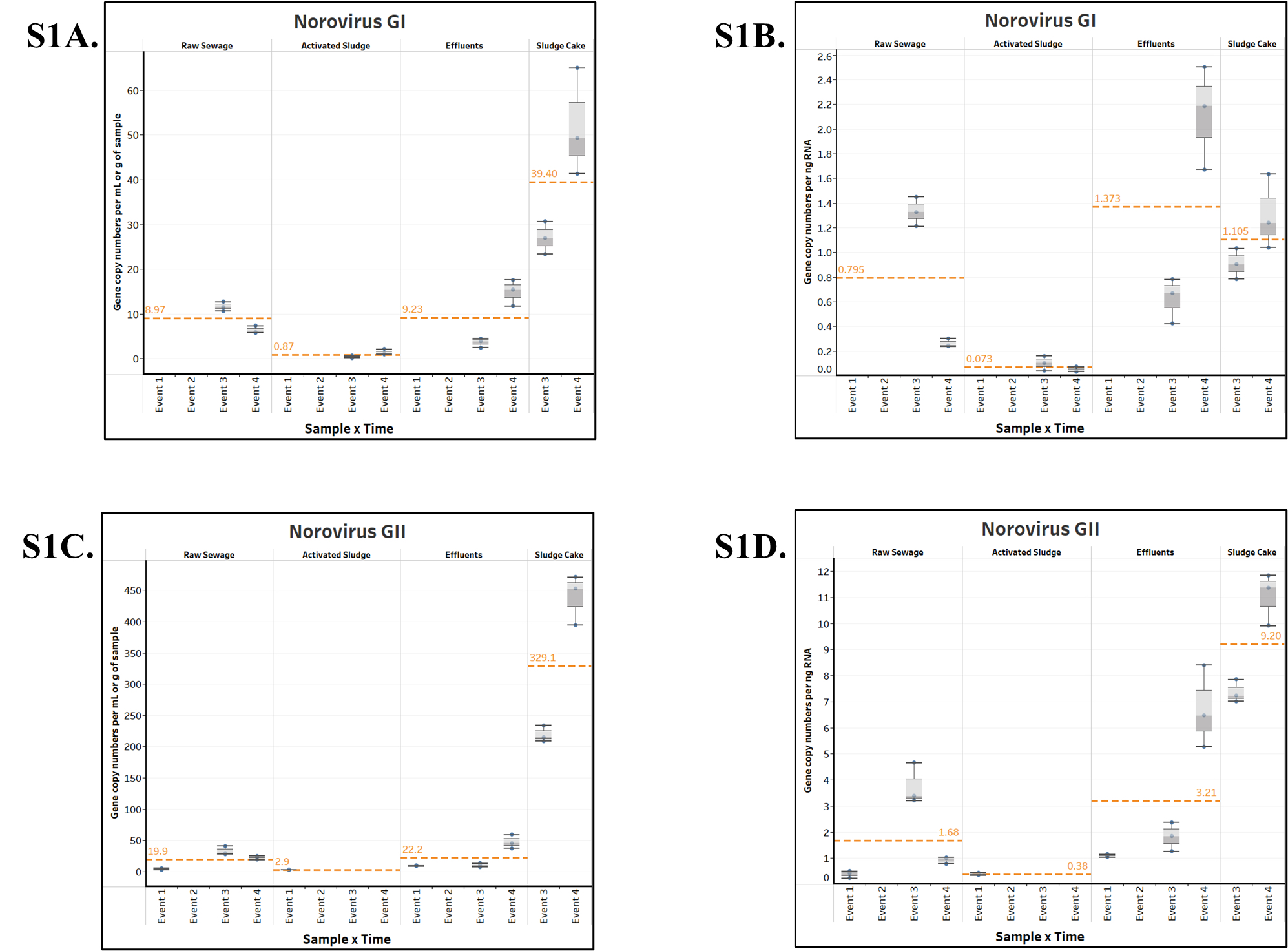

### Figure S2

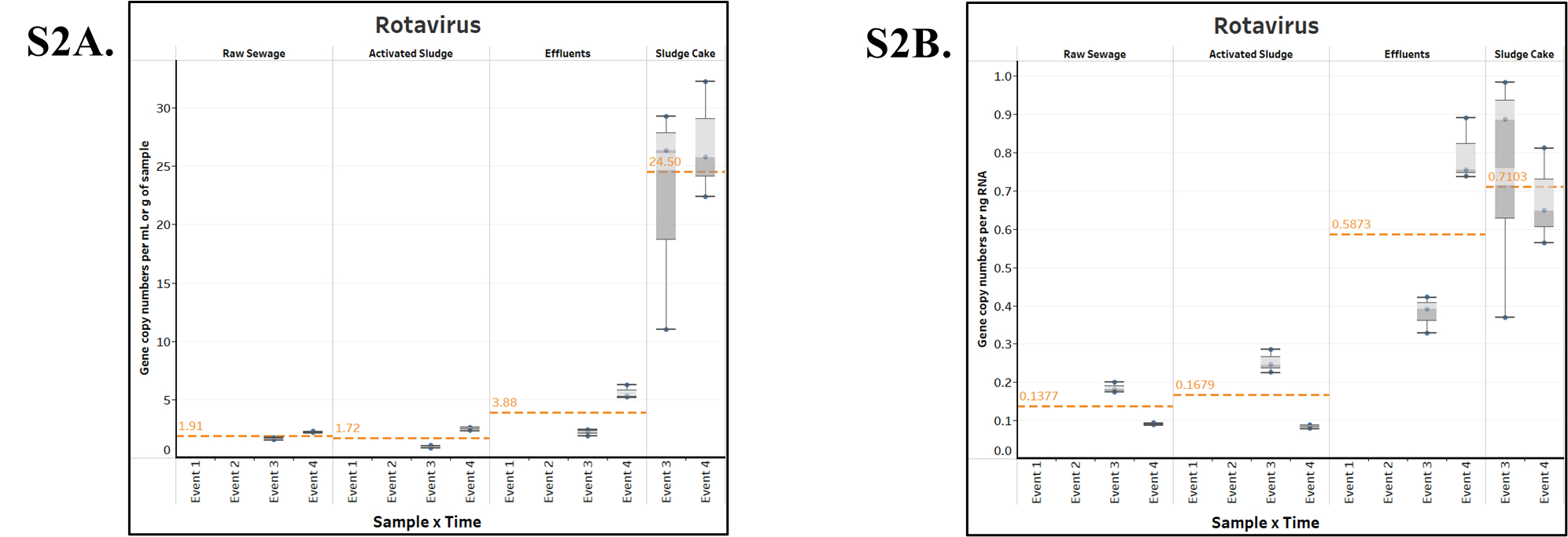

### Figure S3

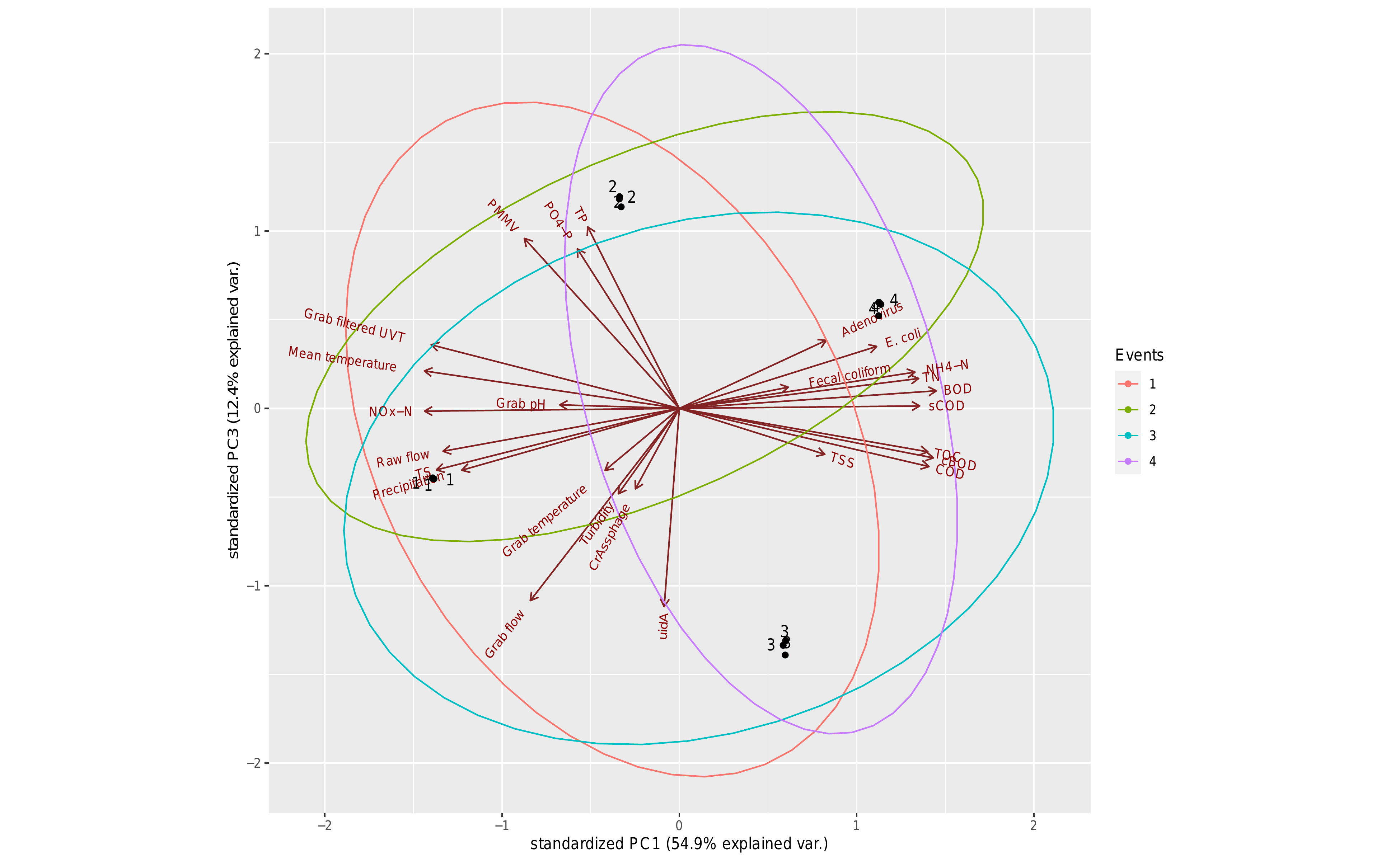

### Figure S4

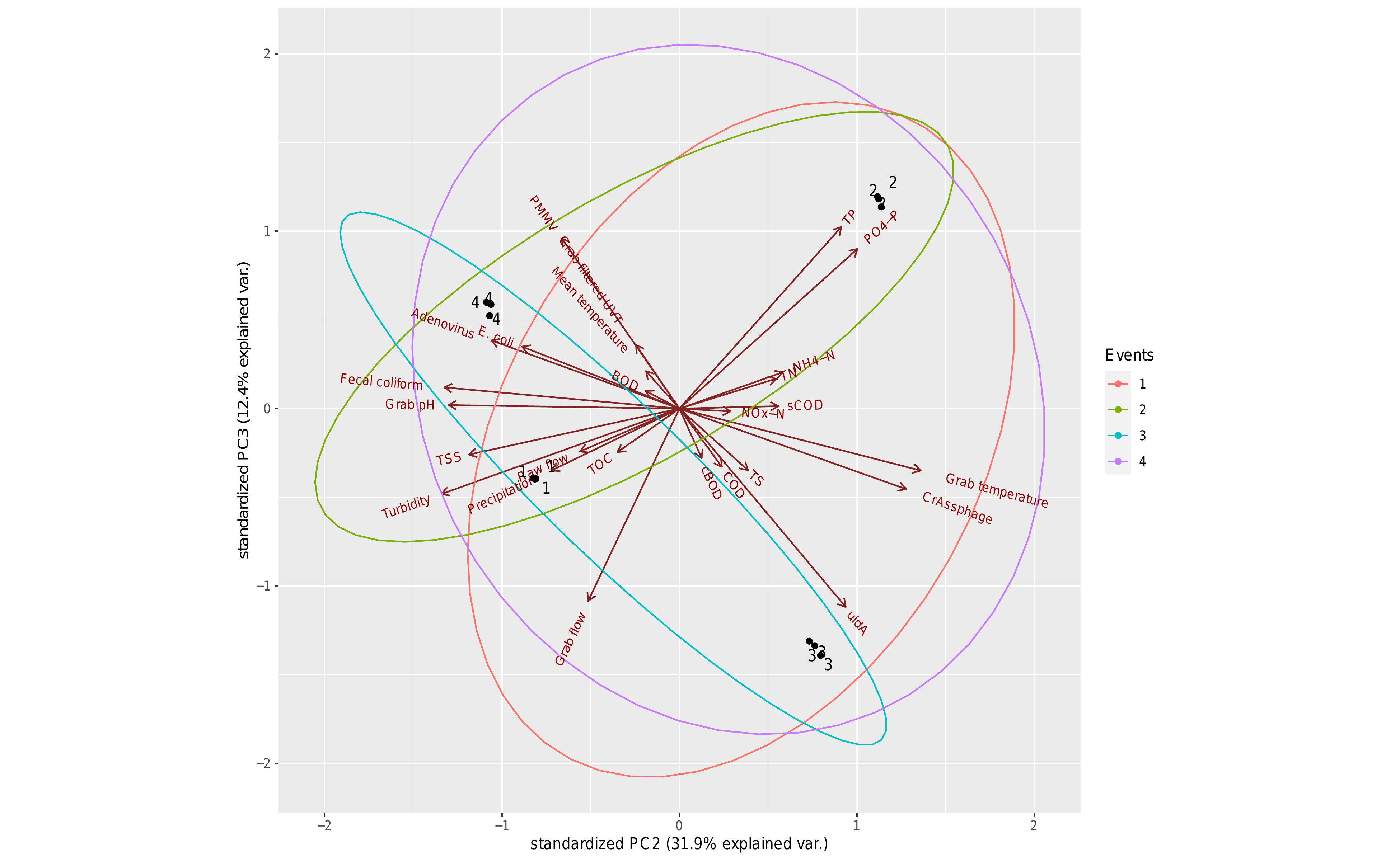
